# Targeted α-Synuclein mRNA Degradation by PMO-Based RNA-Degrading Chimeras

**DOI:** 10.1101/2025.10.17.683180

**Authors:** Ning Wang, Shalakha Hegde, Zhichao Tang, Haiqing Liu, Gang Feng, Lili Niu, Hanyu Li, Kundlik Gadhave, Ramhari Kumbhar, Shu Zhang, Ted M. Dawson, Alexander Pantelyat, Liana S. Rosenthal, Mingyao Ying, Xiaobo Mao, Jingxin Wang

## Abstract

α-Synucleinopathies are devastating neurodegenerative diseases characterized by pathological accumulation of a neuronal protein, α-synuclein (αSyn). Lowering soluble αSyn levels is a promising therapeutic strategy to limit aggregation and neurotoxicity, but directly targeting this protein is hindered by its intrinsically disordered structure and other factors, such as its conformational heterogeneity and intracellular drug delivery barriers. Consequently, increasing attention has been directed toward targeting the *SNCA* transcript, which encodes αSyn. Here, we developed phosphorodiamidate morpholino oligonucleotide (PMO)-based RNA-degrading chimeras (RDCs) that selectively bind the 5′ untranslated region of *SNCA* mRNA and recruit RNase L for targeted RNA degradation. Through the systematic evaluation of 10 RDCs, we identified and optimized 4-D1, which effectively reduced *SNCA* mRNA and αSyn protein expression in HEK293T cells in an RNase L-dependent manner. 4-D1 lowered *SNCA* transcript and αSyn protein levels in both primary cortical neurons from humanized *SNCA* mice and in human induced pluripotent stem cell-derived cortical neurons. This reduction prevented prion-like seeding induced by patient-derived αSyn fibrils and protected neurons from fibril-induced cytotoxicity. Finally, *in vivo* studies confirmed the efficacy of 4-D1 in reducing αSyn mRNA expression in humanized *SNCA* mice. These findings indicate that PMO-based RDCs may represent a promising therapeutic modality for α-synucleinopathies.

**Significance Statement:** Abnormal aggregation of the neuronal protein α-synuclein is central to Parkinson’s disease and related disorders, yet therapeutic candidates that directly target this protein have yet to demonstrate efficacy. in clinical trials. We developed a new strategy that lowers α-synuclein production at the RNA level using phosphorodiamidate morpholino oligonucleotide (PMO)–based RNA-degrading chimeras (RDCs). These molecules recruit a natural RNA-degrading enzyme to selectively destroy the RNA transcript encoding α-synuclein. Our lead RDC reduced α-synuclein levels in cultured cells, humanized mouse and human neurons, blocked α-synuclein pathological aggregation, and protected neurons from toxicity. This study establishes RDCs as a promising therapeutic platform for Parkinson’s disease and other neurodegenerative diseases driven by α-synuclein.

## Introduction

α-Synucleinopathies are a class of devastating neurodegenerative diseases, including Parkinson’s disease (PD), Lewy body dementia, multiple system atrophy, and Alzheimer’s disease with Lewy body pathology (1-3). Despite diverse genetic and environmental contributions (3-6), these disorders share a common histopathological hallmark: the accumulation of misfolded α-synuclein (αSyn) aggregates. Gene dosage effects, such as duplications or triplications of the *SNCA* locus, cause early-onset, autosomal dominant PD (7-9), establishing that αSyn overabundance alone is sufficient to drive α-synucleinopathy. Moreover, αSyn misfolding and aggregation propagate in a prion-like manner, wherein αSyn fibrils seed the conversion of endogenous αSyn monomers and spread between cells, thereby amplifying pathology (3, 6, 10). This process is critically dependent on the presence of endogenous αSyn monomers, as genetic depletion abolishes propagation induced by αSyn preformed fibrils (PFF) (11, 12). Physiologically, αSyn is enriched at presynaptic terminals and implicated in synaptic vesicle trafficking and neurotransmitter release (13, 14). However, *SNCA* knockout studies in animal models reveal only mild phenotypes (15, 16), suggesting that its function is probably partially redundant and compensated by other synucleins. Therefore, reducing αSyn levels represents a rational therapeutic strategy to limit the monomeric substrate required for prion-like propagation.

To date, clinical trials targeting αSyn protein (e.g., prasinezumab and cinpanemab) have yet to demonstrate clinical efficacy. Several factors contribute to these setbacks: (i) αSyn is intrinsically disordered, lacking stable structural epitopes (17, 18), (ii) pathogenic αSyn aggregates primarily form intracellularly and are inaccessible to conventional antibodies (19, 20), and (iii) the existence of diverse αSyn aggregate strains further complicates selective targeting (21, 22). Targeting the *SNCA* transcript upstream of protein synthesis thus provides a tractable strategy to overcome these limitations. RNA-targeting therapeutic modalities, including antisense oligonucleotides (ASOs) (23, 24), RNA interference (RNAi) (23, 25-28), ribozymes (29) and small-molecule translational inhibitors (30, 31), have demonstrated potential to reduce *SNCA* mRNA levels or impair its translation to αSyn. Nevertheless, there remains a critical need for new therapeutic modalities with distinct mechanisms of action, enhanced nuclease stability, and improved target specificity to overcome the limitations of existing RNA-targeting approaches.

Here, we introduce a novel class of phosphorodiamidate morpholino oligomer (PMO)-based RNA-degrading chimeras (RDCs) to lower *SNCA* transcript levels. RDCs exploit endogenous RNase L by conjugating an mRNA-targeting antisense arm to a RNase L-recruiting ligand, converting antisense molecules into active RNA degraders (32). We selected PMOs as the antisense chemistry because they bind RNA targets with high affinity and sequence specificity (33, 34), while exhibiting superior metabolic stability and improved toxicity and immunogenicity profiles (34, 35). Several PMOs have already been approved for clinical use (e.g., eteplirsen, golodirsen) and widely adopted in zebrafish developmental studies for gene knockdown (36, 37). In addition, the neutral backbone of PMOs confers superior solubility in both aqueous and organic solvents, facilitating late-stage chemical modification. By targeting the 5′ untranslated region (UTR) of *SNCA* mRNA, here we report the systematic design and screening of RDCs, demonstrate their RNase L dependence, and validate their ability to reduce both *SNCA* mRNA and αSyn protein levels in primary cortical neurons derived from humanized *SNCA* mice and human induced pluripotent stem cells (iPSC)s. These findings highlight RDC as a promising new therapeutic modality for α-synucleinopathies.

## Results

### Design of PMO-based RDC and screening of RDC binding locations in the *SNCA* 5’ UTR

The RDC modality used in this study is composed of two functional moieties: the RNA-binding moiety and the RNase L recruiter (RLR) moiety (Figure 1A). For RNA-binding, antisense PMOs were employed, which are synthetic oligomers able to bind to RNA through Watson-Crick base pairing using natural nucleobases. Compared to natural RNAs, PMOs feature a backbone composed of phosphorodiamidate linkages and substitute the riboses with morpholino rings (Figure 1B). As a result, PMOs are neutrally charged, a property that not only reduces nonspecific interactions (34) but also makes PMOs more amenable to late-stage chemical modifications. PMOs involved in this study are 17–25 nucleotides in length, as typically used as antisense binders (26), and chemically modified with an alkyne group at the 3’-end, which would be used to conjugate the RLR domain via copper-catalyzed [3+2] cycloaddition (“click chemistry”) (Figure 1B). Previous literature documented a natural RLR, 2’-5’ linked oligoadenylate (2–5A, D2), and its synthetic mimic D1 (Figure 1C), for its application in RNA-degrading chimeras (32, 38, 39). We previously observed that D1 was unexpectedly more active in a small-molecule RDC than natural RLR D2. To screen for optimal RDC-binding locations, we adopted D1 in our initial RDC design, which was fused to each PMO molecule at the 3’ end. At a tolerable location, the RLR domain was tethered with a polyethylene glycol (PEG) linker modified with the azide group, a chemical handle for conjugation reactions with the alkyne group on the PMOs (40).

**Figure 1:**
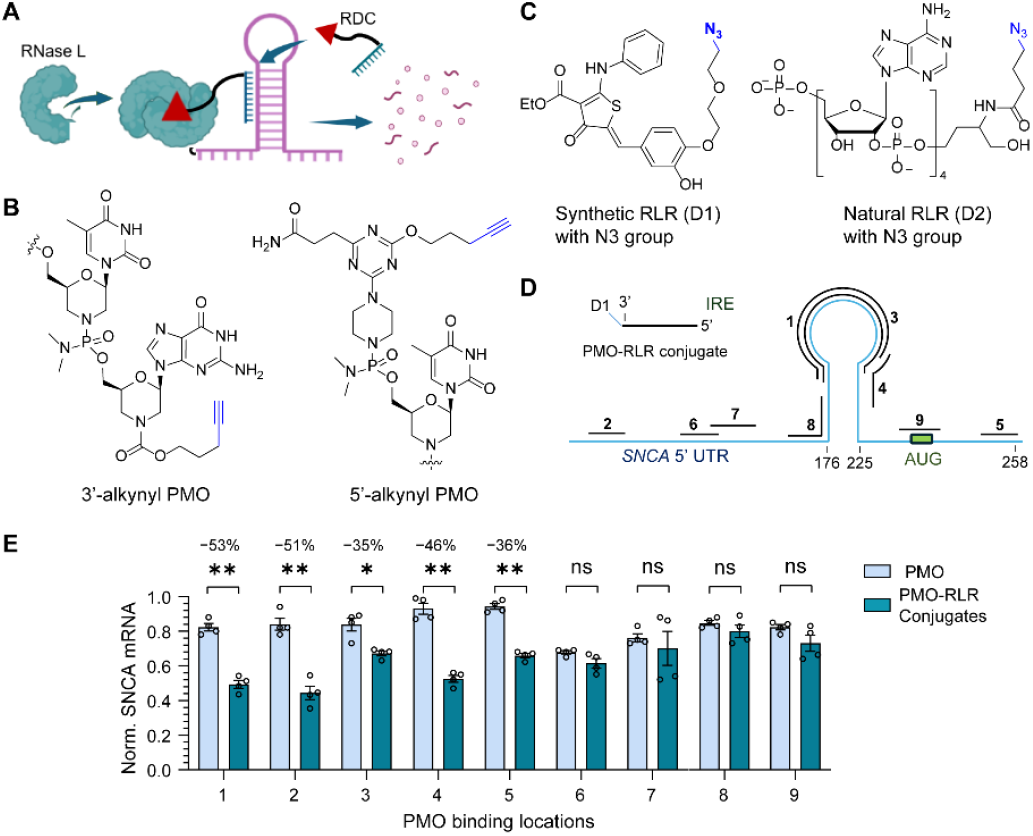
Targeted degradation of *SNCA* mRNA by PMO-RLR conjugates. (A) Schematic of the mechanism of PMO-based RDC targeting the 5’ UTR of *SNCA* mRNA. (B) Representative PMO structures used in this study, including examples of PMOs tagged with a 3’-alkyne (for screening in E) or 5’-alkyne group for conjugation chemistry. (C) Structures of synthetic RLR (D1) and natural 2-5A-based RLR (D2) with an azide (N3) group for conjugation chemistry. (D) Schematic of the *SNCA* 5′ UTR and PMO binding sites, highlighting the IRE loop and the start codon (AUG). The black bars labeled 1–9 represent the binding regions of PMO1 through PMO9, designed to target across the 5′ UTR. (E) Relative *SNCA* mRNA expression levels in HEK293T cells measured using RT-qPCR and normalized to *GAPDH*, following 48-h treatment with PMOs and their corresponding RDCs at 1 μM. Results are shown as mean ± SEM (*n* = 3) relative to the untreated control. The percentages above the bar plots indicate the calculated % of mRNA knockdown for RDCs relative to the corresponding PMOs. Statistical significance is denoted as follows: * *p* < 0.05; ** *p* < 0.01; ns = not significant (*p* > 0.05) using unpaired two-tailed *t*-tests.

We designed a panel of 9 PMOs, targeting across the entire *SNCA* 5’ UTR and the adjacent coding region, with the RDC designated as 1-D1 through 9-D1 (Figure 1D). The design was guided primarily by the feasibility of PMO synthesis and target engagement, with a requirement for G content below 45% and GC content in the range of 25–75% (34). We evaluated the RNA-degrading activity of all the RDCs using reverse transcription-quantitative PCR (RT-qPCR) in HEK293T cells. The activity of each RDC at 1 and 0.1 μM on *SNCA* mRNA level reduction was compared with the corresponding unconjugated PMO (Figure 1E, Supplementary Figure 2). The results indicated that among all the conjugates analyzed, 1-D1, 2-D1, and 4-D1 exhibited the most pronounced knockdown effects of ∼50% reduction on *SNCA* mRNA levels relative to the unconjugated PMOs (Figure 1E). These findings highlight the dependence of binding location for RNA degradation capacity of the RDCs.

### Structural optimization of RDC

To compare the RNA-degrading efficiency of synthetic versus natural RLRs, we synthesized an RDC using PMO1 and the natural RLR D2 (Figure 1C), designated as 1-D2. HEK293T cells were treated with 1-D1 and 1-D2 at a concentration of 1 and 0.1 µM for 48 h, followed by RT-qPCR analysis to measure *SNCA* mRNA levels (Figure S2A). Consistent with our previous results obtained from small-molecule RDCs (41), 1-D1 bearing the synthetic RLR exhibited greater knockdown of *SNCA* mRNA compared to 1-D2, underscoring that the synthetic RLR is also compatible with antisense binders. Interestingly, the synthetic RLR D1 exhibits only micromolar binding affinity for RNase L, which is ∼80,000-fold weaker than the natural ligand D2 (38). The basis for this discrepancy between binding affinity and functional activity remains unclear.

Next, we sought to investigate whether the site of RLR conjugation on the PMO impacts the RDC activity. Specifically, we tested whether conjugating the RLR (D1) to either the 3′ or 5′ end of the PMO would affect *SNCA* mRNA reduction in cells. To this end, we synthesized an RDC 4-5’-D1 by using an alkyne group at the 5′ end of PMO4 for D1 conjugation (Figure 1B). HEK293T cells were treated with 4-5’-D1 and compared with 4-D1 (3’-conjugation) at 1 and 0.1 μM. Interestingly, both conjugates achieved similar levels of *SNCA* knockdown across the tested concentrations (Figure S2B), suggesting that the conjugation termini of the RLR on the PMO backbone do not significantly influence the activity of the RDC in cells.

### Evaluation of dose–response and time-dependent activity of RDCs

To gain insight into the RDC activity and kinetics, we performed a dose response and time-course analysis for two active RDCs, 4-D1 and 2-D1. HEK293T cells were treated with increasing concentrations of 4-D1 and 2-D1 ranging from 0.03 to 1.5 µM. Both RDCs induced a robust, concentration-dependent reduction of *SNCA* mRNA, with half-maximal effective concentrations (EC_50_) of ∼100 nM (Figure 2A). We next evaluated the effects of 4-D1 treatment at 1 µM and 0.1 µM over a 4-day period. RNA knockdown was detectable as early as 24 h post-treatment and reached plateau by Day 3 (Figure 2B).

**Figure 2:**
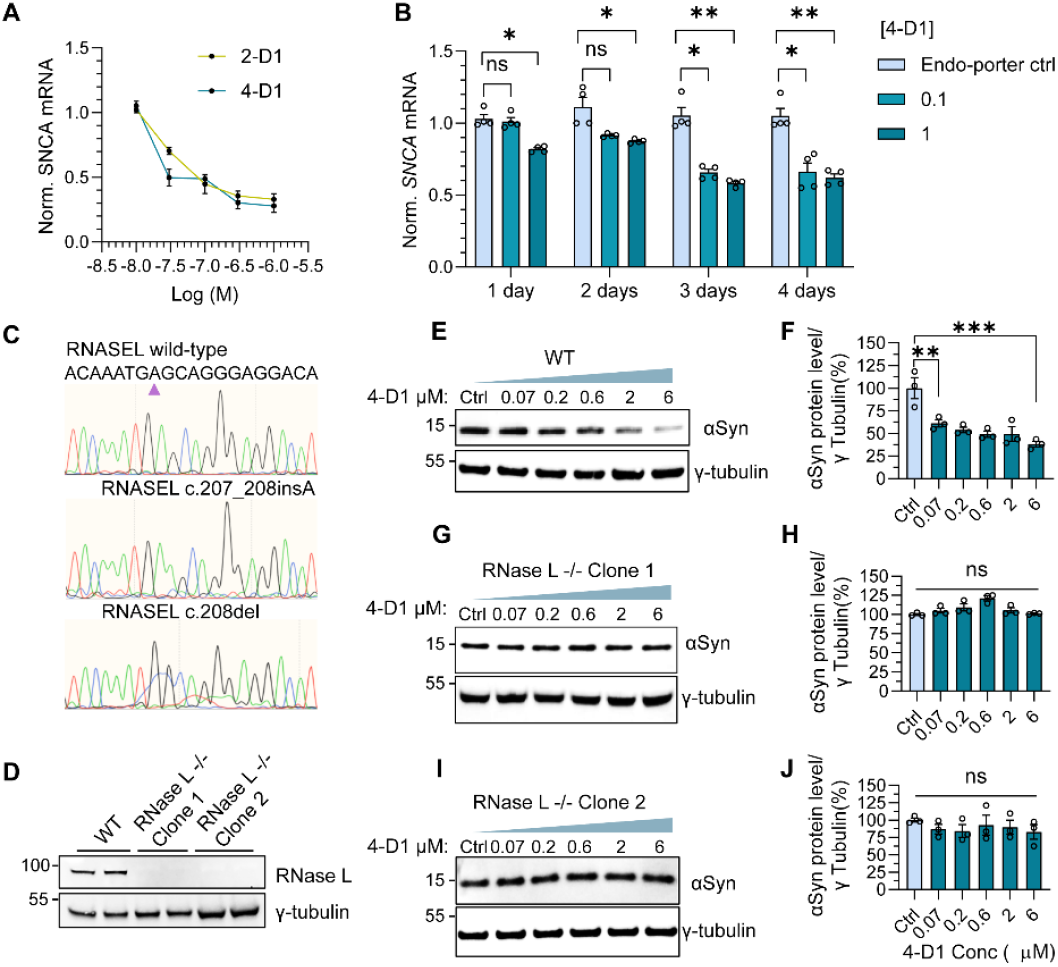
Profiles of RDC activity: dose-response, time-course, and mechanistic evaluation. (A) Dose response curves of *SNCA* mRNA levels in HEK293T cells measured by RT-qPCR following 48-h treatment with various concentrations of 2-D1 and 4-D1. mRNA levels were normalized to the *GAPDH* mRNA level. Data points are shown as mean ± SEM (*n* = 3), relative to the control (cells treated with Endo-Porter alone). (B) Time-course analysis of *SNCA* mRNA knockdown in HEK293T cells treated with 1 μM and 0.1 μM 4-D1. mRNA levels were normalized to β-actin and are presented as mean ± SEM (*n* = 4) relative to the control. The statistical significance was evaluated via one-way ANOVA with Tukey’s multiple comparisons test (ns, *p* > 0.05, \**p* < 0.05, \*\**p* < 0.01). (C) Sanger sequencing chromatograms of the CRISPR-cas9-targeted region in the RNASEL gene from wild-type HEK293T cells (top) and two RNase L^−/−^ clones (middle and bottom). (D) Immunoblotting with anti-RNase L antibody confirmed loss of RNase L expression in both RNase L^−/−^ clones, with γ-tubulin as a loading control. (E) Representative Western blot analysis of αSyn levels in HEK293T cells following treatment with various concentrations of 4-D1 (0.07–6 μM, 48 h), with γ-tubulin as a loading control. (F). Densitometric quantification of αSyn levels after immunoblotting, normalized to γ-tubulin. Data represents the mean of three biological replicates. (G–J) Representative western blot analysis of αSyn levels in wild-type and RNase L^−/−^ cells following the treatment with varied concentrations of 4-D1 (0.07–6 µM, 48 h) (G, I). Quantification of αSyn protein levels (normalized to γ-tubulin) relative to the control samples (H, J) (*n* = 3). The statistical significance was evaluated via one-way ANOVA with Tukey’s multiple comparisons test (ns, *p* > 0.05, \**p* < 0.05, \*\**p* < 0.01, \*\*\**p* < 0.001).

### RDCs reduced monomeric αSyn protein level in an RNase L-dependent manner

To further investigate the mechanism underlying RDC-mediated αSyn reduction, we generated an RNase L knockout (RNase L^−/−^) HEK293T cells using CRISPR-Cas9. We selected two colonies with different homozygous frameshift mutations of RNase L (Figure 2C), and immunoblot analysis confirmed the knockout by showing loss of RNase L expression in both clones (Figure 2D). In wild-type HEK293T cells, 4-D1 induced a dose-dependent reduction of αSyn protein expression, achieving ∼50% reduction at 6 µM compared to untreated control (Figure 2B). In contrast, treatment of the RNase L^−/−^ cells with various concentrations of 4-D1 abolished αSyn knockdown activity in both clones (Figure 2G–2J). These results underscore the RNase L-dependence in mediating the activity of the RDCs.

### RDC 4-D1 reduces αSyn levels and prevents pathological seeding by PD αSyn fibrils in humanized *SNCA* mouse neurons

To extend our cell line findings to a disease-relevant context, we evaluated 4-D1 in primary cortical neurons derived from the humanized *SNCA* PAC-Tg (*SNCA*_WT_) mouse model (Jackson Laboratory, Stock No: 010710) (42, 43), which carries the full-length human wild-type *SNCA* gene including its UTRs on a murine *Snca*-null background (Fig. 3A). 4-D1 treatment produced robust target knockdown, with RT-qPCR analysis revealing an ∼50% reduction in *SNCA* mRNA levels relative to control neurons (Fig. 3B).

**Figure 3.**
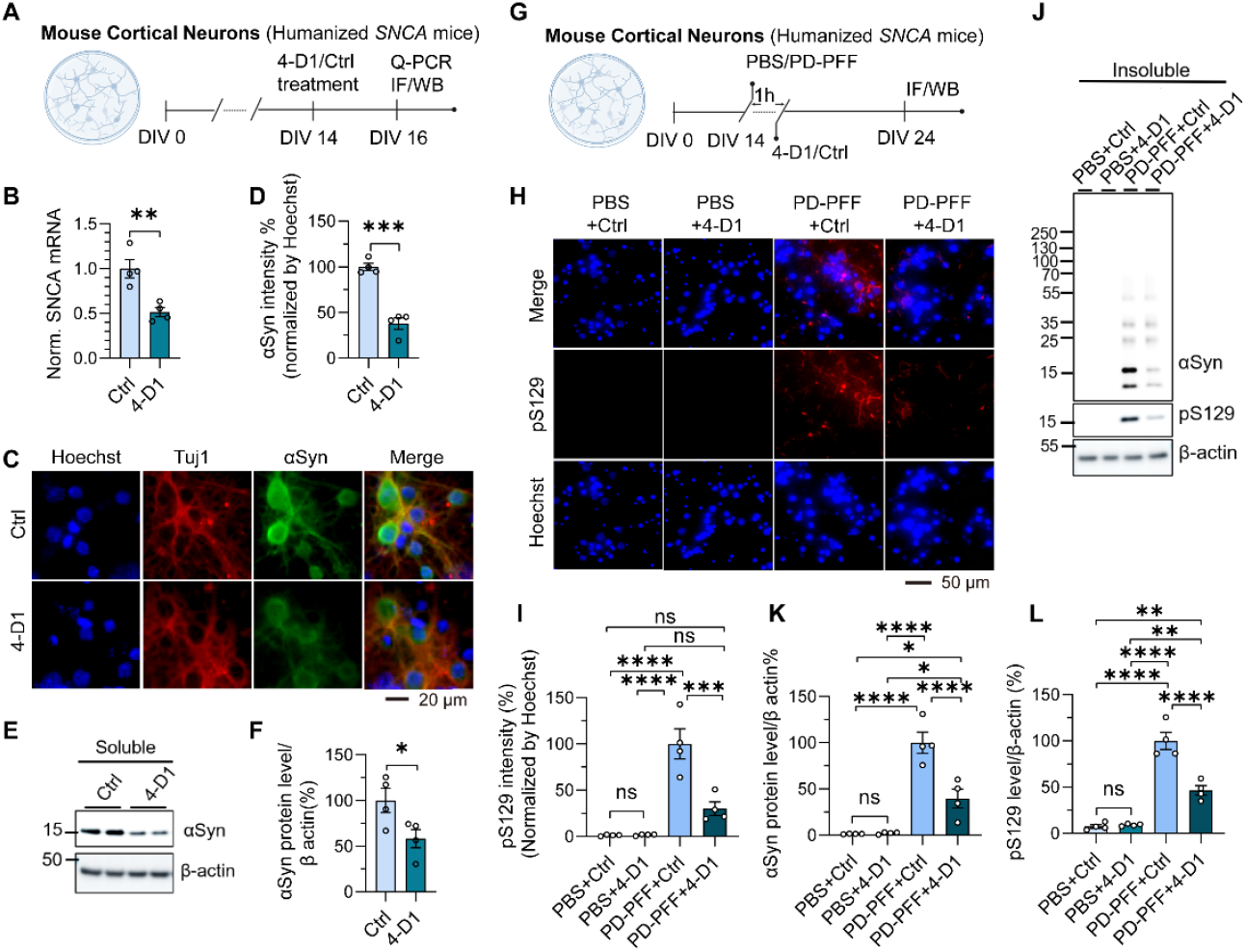
4-D1 reduced monomeric αSyn protein level and pS129 level in cultured humanized *SNCA* neurons. (A) Schematic of the experiment design of 4-D1 or unconjugated PMO4 (Ctrl) in humanized *SNCA* neurons. (B) *SNCA* mRNA level was decreased in 4-D1-treated humanized *SNCA* neurons compared to the Ctrl group (*n* = 4). (C) Immunofluorescence analysis of Tuj1 (red), αSyn (green), and Hoechst (blue) in humanized *SNCA* neurons. Scale bar: 20 μm. (D) Quantitative results of αSyn intensity of humanized *SNCA* neurons with Ctrl or 4-D1 treatment (*n* = 4). (E, F) SDS–PAGE data showed αSyn level with 4-D1 or Ctrl treatment (*n* = 4). (G) Schematic of the experiment design of PD-PFF treatment following 4-D1 or Ctrl in humanized *SNCA* neurons. (H) Immunofluorescence analysis of αSyn phosphorylated at serine 129 (pS129, red) and Hoechst (blue) in humanized *SNCA* neurons. Scale bar: 50 μm. Values are expressed as mean ± SEM. \**p* < 0.05, \*\**p* < 0.01, \*\*\**p* < 0.001, unpaired two-tailed Student’s t test. (I) Quantitative results of H (*n* = 4). (J–L) αSyn protein level (K) and pS129 level (L) with PD-PFF treatment following 4-D1 or Ctrl treatment in humanized *SNCA* neurons (*n* = 4). The statistical significance was evaluated via one-way ANOVA with Tukey’s multiple comparisons test (ns, *p* > 0.05, \**p* < 0.05, \*\**p* < 0.01, \*\*\**p* < 0.001. \*\*\*\**p* < 0.0001).

This reduction in transcript abundance was accompanied by a parallel decrease in αSyn protein. Immunofluorescence analysis showed a marked reduction in αSyn staining, with quantification confirming an ∼50% decrease in total protein levels (Fig. 3C, D). Immunoblotting of the Triton X-100-soluble fraction further corroborated these findings, demonstrating an ∼50% decrease in soluble αSyn with 4-D1 treatment (Fig. 3E, F).

Because a key pathogenic mechanism in α-synucleinopathies involves prion-like propagation of misfolded αSyn, we next tested whether lowering endogenous αSyn could protect against seeding by pathogenic fibrils. Humanized *SNCA* neurons were pretreated with 4-D1 or unconjugated PMO4 (Ctrl) prior to exposure to αSyn preformed fibrils (PFF) amplified from the cerebrospinal fluid (CSF) of a patient with Parkinson’s disease (PD-PFF) (Fig. 3G) (22). The fibrillar morphology of PD-PFF was confirmed by transmission electron microscopy before and after sonication, which fragmented fibrils into seeding-competent species (Fig. S3). Pathology induction was monitored by αSyn phosphorylated at serine 129 (pS129), a hallmark of pathological αSyn aggregates (19, 21, 22, 44-48).

Pre-treatment with 4-D1 significantly attenuated PD-PFF-induced pS129-positive αSyn pathology, with quantitative immunofluorescence showing a ∼75% reduction in the pS129 signal (Fig. 3H, I). To confirm this result biochemically, we analyzed Triton X-100-insoluble fractions from neuronal lysates. Immunoblotting demonstrated a ∼70% decrease in insoluble αSyn (Fig. 3J, K) and an accompanying ∼50% reduction in insoluble pS129 signal (Fig. 3J, L) in 4-D1-treated cultures compared with controls.

> *Collectively, these data demonstrate that 4-D1 not only effectively lowers SNCA mRNA and αSyn protein, but also markedly attenuates the seeding and pathological conversion induced by disease-derived αSyn fibrils in a primary neuron model*.

### RDC 4-D1 reduces αSyn levels and prevents pathological seeding by PD αSyn fibrils in human iPSC-derived neurons

To validate these findings in a fully human neuronal system and rule out species-specific artifacts, we used cortical glutamatergic neurons differentiated from human iPSCs *via* a widely adopted Ngn2-driven protocol (49), which yields a highly pure excitatory neuronal population (Fig. S4). As shown in the experimental schematic (Fig. 4A), neurons were treated with 4-D1 or Ctrl, followed by assessment of *SNCA* mRNA and αSyn protein. 4-D1 treatment produced a significant ∼40% reduction in *SNCA* transcript levels as determined by RT-qPCR (Fig. 4B). Immunofluorescence analysis confirmed that this transcript knockdown translated into protein suppression (Fig. 4C), with quantitative analysis revealing a ∼75% reduction in αSyn intensity in both soma (Fig. 4D, F) and neurites (Fig. 4E, G) of 4-D1-treated neurons.

**Figure 4.**
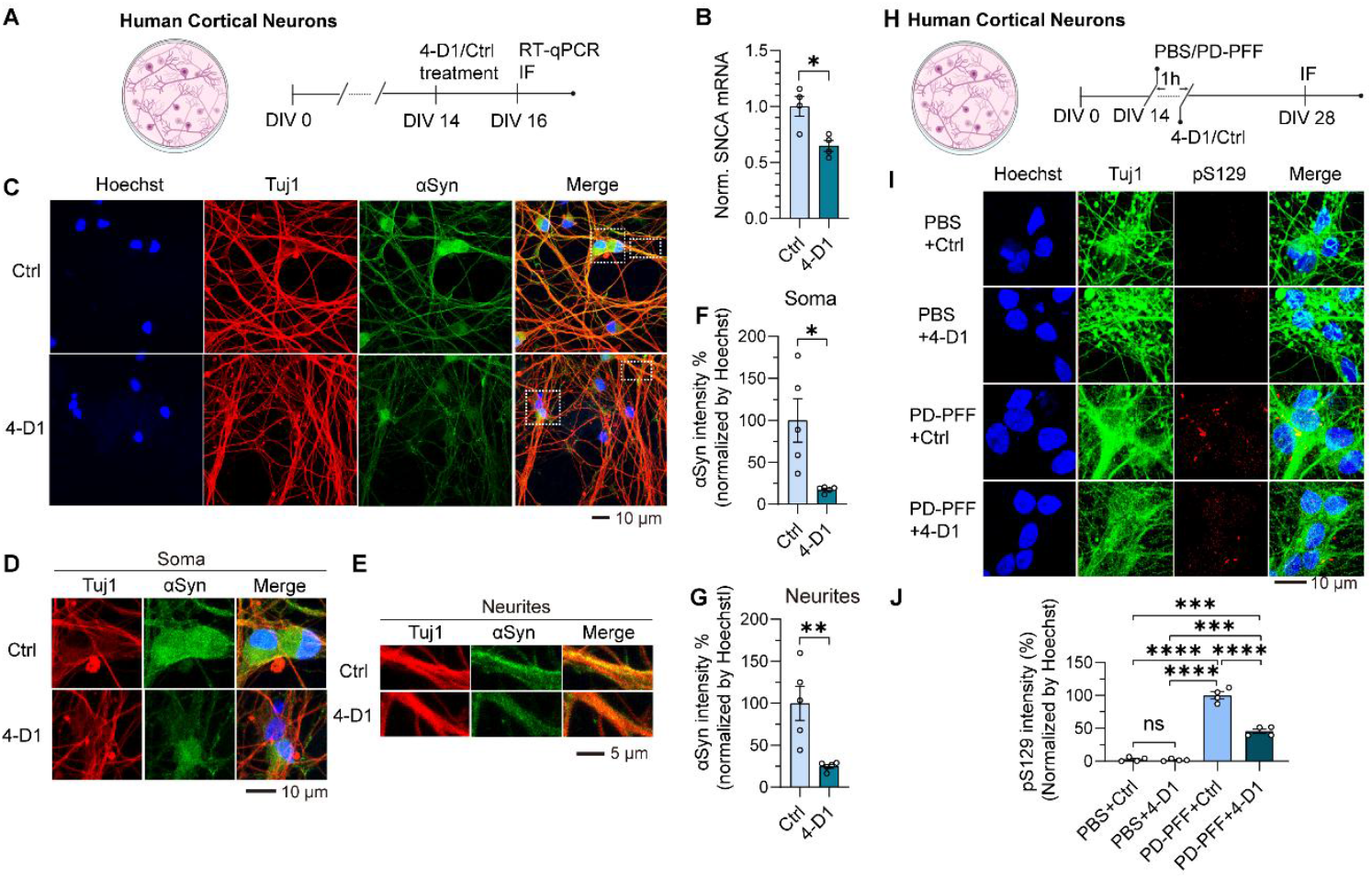
4-D1 reduced monomeric αSyn protein level and pS129 level in human neurons. (A) Schematic of the experiment design of 4-D1 or unconjugated PMO4 (Ctrl) in human cortical neurons. (B) *SNCA* mRNA level was decreased in 4-D1-treated human neurons compared to the Ctrl group (*n* = 4). (C) Immunofluorescence analysis of Tuj1 (red), αSyn (green) and Hoechst (blue) in human neurons. Scale bar: 10 μm. (D, E) Zoomed images of soma (D) and neurites (E) of human neurons with Ctrl or 4-D1 treatment. (F, G) Quantitative results of αSyn intensity in soma (F) and neurites (G) of human neurons with Ctrl or 4-D1 treatment (*n* = 4). Values are expressed as mean ± SEM. \**p* < 0.05, \*\**p* < 0.01, unpaired two-tailed Student’s t test. (H) Schematic of the experiment design of PD-PFF treatment following 4-D1 or Ctrl in human cortical neurons. (I) Immunofluorescence analysis of Tuj1 (green), pS129 (red), and Hoechst (blue) in human neurons. Scale bar: 10 μm. (J) Quantitative results of I (*n* = 4). The statistical significance was evaluated via one-way ANOVA with Tukey’s multiple comparisons test (ns, *p* > 0.05, \**p* < 0.05, \*\**p* < 0.01, \*\*\**p* < 0.001. \*\*\*\**p* < 0.0001).

We next asked whether 4-D1 could protect these human neurons from pathological seeding by PD-PFF. Two weeks after exposure, Ctrl-treated neurons developed pS129-positive inclusions, whereas pre-treatment with 4-D1 markedly suppressed pathology (Fig. 4H, I). Quantitative analysis showed a ∼50% decrease in pS129 immunoreactivity compared to controls cultures (Fig. 4J).

> *Collectively, these results confirm the efficacy of 4-D1 in a human neuronal context, demonstrating its ability to lower physiological αSyn levels across neurons and attenuate the downstream pathological conversion induced by disease-derived αSyn fibrils*.

### RDC 4-D1 rescues humanized *SNCA* neurons from toxicity induced by PD αSyn fibrils

Having established that 4-D1 reduces αSyn pathology, we next asked whether this translated into a functional benefit by preventing neuronal death. In the humanized *SNCA* primary neuron model (Fig. 5A), exposure to PD-PFF induced neurotoxicity in cultures treated with the Ctrl, resulting in the loss of ∼50% of neurons as quantified by NeuN immunostaining (Fig. 5B, C). Similarly, human iPSC-derived cortical neurons exposed to PD-PFF for three weeks exhibited marked neurotoxicity (Fig. 5D, E).

**Figure 5.**
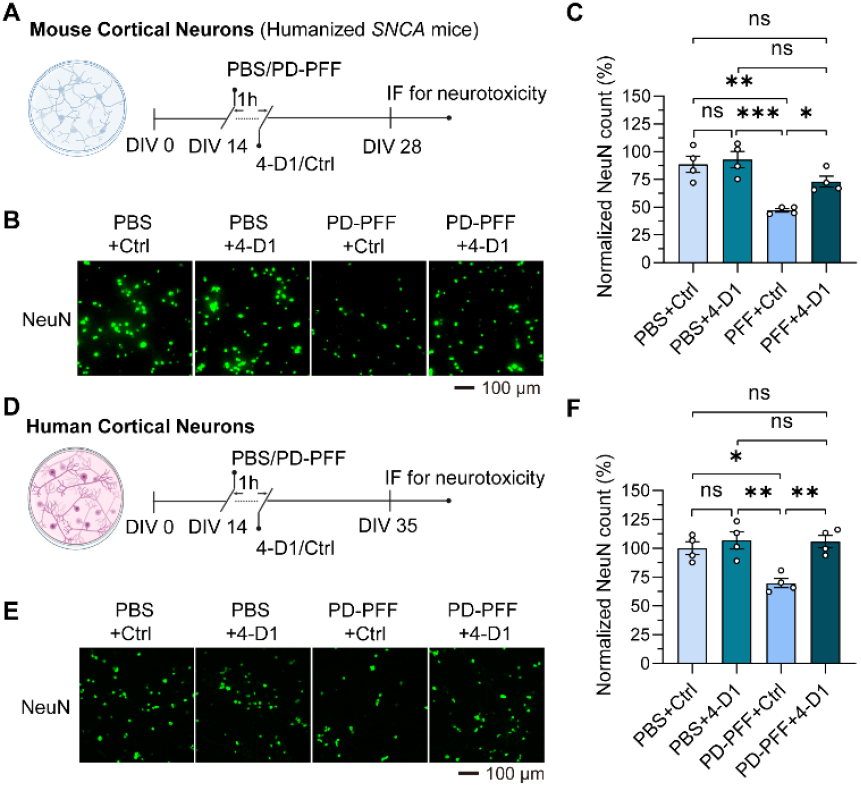
4-D1 alleviated PD-PFF-induced neurotoxicity in humanized *SNCA* neurons and human neurons. (A) Schematic of the experiment design of 4-D1 or unconjugated PMO4 (Ctrl) treatment in primary cortical neurons derived from humanized *SNCA* mice. (B, C) Neurotoxicity assessment with anti-NeuN immunostaining and quantification in primary cultured neurons with 10 μg/mL amplified PD-PFF (*n* = 4). scale bar, 100 µm. (D) Schematic of the experiment design of 4-D1 or control treatment in human cortical neurons. (E, F) Neurotoxicity assessment with anti-NeuN immunostaining and quantification in primary cultured neurons with 10 μg/mL amplified PD-PFF (*n* = 4). The statistical significance was evaluated via one-way ANOVA with Tukey’s multiple comparisons test (ns, *p* > 0.05, \**p* < 0.05, \*\**p* < 0.01, \*\*\**p* < 0.001).

Pre-treatment with 4-D1 conferred substantial neuroprotection. While Ctrl-treated cultures retained 50% neuronal viability after PD-PFF administration, 4-D1-treated cultures showed restored viability of 75%. This indicates that 4-D1 treatment rescued approximately half of the neuronal loss in humanized *SNCA* primary neurons (Fig. 5B, C) and completely preserved human cortical neurons (Fig. 5E, F). These findings highlight the therapeutic potential of 4-D1, which not only halts αSyn pathology propagation but also preserves neuronal viability under disease-relevant conditions.

### Direct cortical administration of RDC 4-D1 reduces *SNCA* mRNA levels *in vivo*

To determine if 4-D1 could engage its target *in vivo*, we performed a single stereotaxic injection of 4-D1 or Ctrl into the motor cortex of humanized *SNCA* mice (Fig. S5A). Five days post-injection, motor cortex tissue was collected for analysis. Quantitative PCR revealed that 4-D1 treatment produced a ∼50% reduction in human *SNCA* mRNA compared with the hemisphere receiving the Ctrl (Fig. S5B). *These results provide clear in vivo evidence of target engagement, demonstrating that 4-D1 is active and capable of mediating mRNA degradation within the brain*.

## Discussion

During mechanistic studies of αSyn-lowering RDCs, we initially hypothesized that a 5′-UTR– targeting PMO might also exert inhibitory activity by sterically hindering ribosome loading and thereby synergizing with mRNA degradation. However, our RNase L knockout experiments demonstrated that PMO alone was inactive in reducing αSyn expression, indicating that αSyn-lowering RDCs in this study depends almost entirely on RNase L–mediated mRNA knockdown. Nonetheless, for other transcripts, particularly those whose translation initiation is more vulnerable to steric blockade, a synergistic effect of ribosome interference combined with targeted mRNA degradation could be realized in RDCs, potentially enhancing potency and broadening the therapeutic scope of this pharmacological modality.

RDCs exploit endogenous RNase L by conjugating an antisense “guide” arm to an RNase L– recruiting ligand (“effector” arm), thereby converting otherwise inactive RNA-binding molecules (e.g., PMOs alone in this study) into active RNA degraders (50). This concept was first demonstrated by the Silverman group in 1993 (32). In 2018, the Disney group extended this strategy by tethering a synthetic RNase L–binding moiety to a small-molecule RNA ligand (51). Recently, an RDC termed Syn-RiboTAC was reported, in which a small-molecule RNA binder targets the iron-responsive element (IRE) in the 5′ untranslated region (UTR) of *SNCA* mRNA, showing promising activity at micromolar concentrations (30). Compared with this small-molecule– based RDC approach, our PMO-based RDCs exhibit higher potency, highlighting the advantages of antisense-guided recruitment of RNase L for selective RNA degradation. However, our current PMO-based RDCs are not inherently cell-permeating in cell culture, and future work will focus on developing delivery strategies to enable their therapeutic applications.

In our *in vivo* studies, we did not observe a corresponding reduction in αSyn protein levels in the mouse brain despite clear knockdown of *SNCA* mRNA. This may be explained by the characteristics of the PAC-Tg (*SNCA*_WT_) mouse model, which exhibits a large, ∼50-fold overexpression of *SNCA* transcript compared to wild-type mouse, but nearly no increase in αSyn protein (see mouse strain description at the Jackson Laboratory, Stock No. 010710). This disconnect suggests the presence of potent post-transcriptional or translational regulatory mechanisms that buffer αSyn protein expression, limiting the dynamic range within which RNA knockdown can alter protein output. Consequently, this model may underestimate the capacity of targeted RNA degradation approaches to lower αSyn protein. Future studies in more physiologically relevant models that better recapitulate *SNCA* expression and αSyn homeostasis will be critical to fully assess the therapeutic potential of RDCs.

In conclusion, our study demonstrates that PMO-based RDCs effectively reduce *SNCA* mRNA and αSyn protein by targeting specific 5’ UTR sequences. Conjugation of PMOs with the synthetic RNase L-recruiting ligand D1 was shown to be more efficacious than with natural ligand D2. Comparable activity from 3’ and 5’ end PMO conjugation indicates flexibility in RDC design. Notably, the absence of activity in RNase L knockout cells confirms that the observed effects are mediated by RNase L-dependent RNA degradation.

Given that αSyn fibrils derived from Lewy body diseases exhibit structural and functional differences from mouse αSyn PFF (52), we employed PD patient-derived αSyn strains in both humanized *SNCA* primary cortical neurons and iPSC-derived human cortical neurons. The demonstrated efficacy of 4-D1 in reducing αSyn monomer levels and inhibiting patient-derived seed-induced pathology propagation and neurotoxicity provides strong rationale for subsequent *in vivo* validation studies. Collectively, these findings highlight the potential of PMO-based RDCs as a targeted therapeutic approach for disorders characterized by αSyn aggregation, including PD and other α-synucleinopathies.

## Materials and Methods

### General Procedure for Synthesizing PMO-based RDCs

A mixture of 3’-alkyne-modified *SNCA* PMO (1 mM in water, 50 µL; see **Supplementary Table 1** for sequences and modification annotations), D1-N_3_ (10 mM in DMSO, 15 µL, 3 eq; synthesized as described in **Supplementary Methods**), or D2-N_3_ (10 mM in water, 15 µL, 3 eq), CuSO_4_ (100 mM in water, 5 µL, 10 eq), THPTA (100 mM in water, 5 µL, 10 eq), and sodium ascorbate (100 mM in water, 5 µL, 10 eq) in 370 µL deionized water and 50 µL DMSO was stirred at room temperature overnight under nitrogen. LC-MS analysis confirmed completion of the reaction. The mixture was filtered through a 0.22 µm membrane and purified by reverse-phase HPLC (2–98% acetonitrile gradient in 0.1% ammonium formate aqueous solution). The desired fractions were collected, lyophilized, and reconstituted in molecular biology-grade water at 1 mM as stock solution (**Supplementary Methods**).

### RDC treatment and Quantitative Reverse Transcription PCR (RT-qPCR) Assay

The wild-type HEK293T cells were seeded at 3 × 10^5^ cells/ml in 12-well plates in 1 mL growth medium (10% FBS in DMEM medium) at 37 °C for 3 h. The cells were then treated with the PMOs or PMO-based RDCs at various concentrations for different time points in the presence of 0.3% Endo-Porter PEG (Gene Tools, OT-EP-PEG-1). During 4-day treatments, the concentration of Endo-Porter was reduced from 0.3% to 0.2%. After treatment, the supernatant was aspirated from each well and the total RNA was then extracted from the cells using RNeasy Mini kit (Qiagen, 74104). The total RNAs were quantified by ultraviolet absorption at 260 nm (NanoDrop 1000, Thermo Fisher, Waltham, MA, USA). cDNAs were synthesized from 500 ng of total RNA for each sample using M-MLV reverse transcriptase (Promega, M1701) and (dT)_25_ according to the manufacturer’s protocol. 1 µl of cDNA mixture was used in a 15 µl RT-qPCR reaction (SYBR Green, Apex-Bio, K1070). The human *GAPDH* mRNA level was used as the reference for normalization. RT-qPCR primer sequences:

*SNCA*-FW: 5’-CAGAAGGGGCCCAAGAGAGG; *SNCA*-RV:5’ CCTCCTTGGTTTTGGAGCCT GAPDH-FW: 5’-GACAAGGCTGGGGCTCATTT; GAPDH-RV: 5’-CAGGACGCATTGCTGATGAT

### Immunoblotting

Wild-type or RNase L^−^/^−^ HEK293T cells were seeded at a density of 3 × 10^5^ cells/ml in 12-well plates with 1 mL of growth medium and incubated at 37 °C for 3 h. Cells were then treated with RDCs as described above. At 48 h post-treatment, cells were washed once with 1× PBS, harvested, and lysed in 1× RIPA buffer (Cell Signaling Technology, 9806). Lysates were vortexed for 1 min and centrifuged at 17,000 × g for 10 min at 4 °C to pellet debris. The supernatant was collected, mixed with an equal volume of 2× SDS sample buffer, and heated at 95 °C for 5 min. Neuron samples collected from humanized *SNCA* mice were homogenized with TX-soluble buffer (50 mM Tris [pH 8.0], 150 mM NaCl, 1% Triton X-100) containing protease and phosphatase inhibitors (Sigma-Aldrich, 11697498001). The supernatants were collected for soluble fraction after centrifugation (20,000 × g, 20 min), and the pellets were resuspended in TX-insoluble buffer (containing 2% SDS) with protease and phosphatase inhibitors. Proteins were resolved by SDS-PAGE using a 4–12% Bis-Tris gel (Invitrogen, NW04125BOX) and transferred to a PVDF membrane (Invitrogen, IB34002). Membranes were fixed in 0.04% paraformaldehyde in 1× PBS for 30 min at room temperature to preserve low-molecular-weight proteins, followed by standard immunoblotting. Following protein transfer, membranes were blocked with 5% bovine serum albumin (BSA) in Tris-buffered saline with 0.1% Tween-20 (TBST) for 1 hour at room temperature. Membranes were then incubated overnight at 4 °C with primary antibodies diluted in 5% BSA buffer, targeting RNase L (1:500, Cell Signaling Technology, 27281), αSyn (1: 1000, BD Biosciences, 610786), phosphorylated S129 (pS129) αSyn (1:1000, Abcam, ab51253), γ-tubulin (1:5000, Sigma-Aldrich, T6557), or β-actin (1: 5000, Sigma-Aldrich, A3854). After washing with TBST, membranes were incubated with an HRP-conjugated anti-mouse or anti-rabbit IgG secondary antibody diluted 1:1000-1:2000 in 5% BSA buffer (Cell Signaling Technology, 7076S) for 1 h at room temperature. Protein bands were visualized using the SuperSignal™ West Pico PLUS chemiluminescent substrate (Thermo Fisher, 34577) and imaged on an iBright FL-1500 Imaging System (Thermo Fisher, Waltham, MA, USA). Band intensities were quantified using iBright Analysis Software, and αSyn levels were normalized to γ-tubulin or β-actin.

### Animal procedures

All animal procedures were rigorously reviewed and approved by the Johns Hopkins Animal Care and Use Committee (IACUC), ensuring that all experiments were conducted ethically and in strict accordance with the NIH Guide for the Care and Use of Laboratory Animals. For this study, we used humanized wild-type α-synuclein (Humanized *SNCA*) mice (FVB;129S6-Snca^tm1Nbm^ Tg (SNCA)1Nbm/J, Strain #010710), which were originally obtained from The Jackson Laboratory (Bar Harbor, ME, USA). Mice were housed in ventilated cages under a controlled 12-hour light/dark cycle and were provided with ad libitum (free) access to standard chow and water throughout the duration of the experiments.

### Amplification of pathogenic αSyn with seeding amplification assay (SAA) and human PD-PFF preparation

Pathological αSyn seeding activity was quantified using an SAA protocol adapted from our previous work, employing a Qsonica microplate horn sonicator system (53). Recombinant αSyn monomer was first cleared of aggregates by ultracentrifugation (100,000 × g, 30 min, 4°C, Beckman Coulter, A95761). The reaction mixture, consisting of 0.3 mg/mL αSyn substrate in SAA buffer (1% Triton X-100 in PBS) and silicon beads, was seeded with 10 µL of cerebrospinal fluid (CSF) from PD patients. The reaction plates were subjected to repeated cycles of sonication (25% amplitude, 30 sec) and incubation (37°C, 29.5 min) for 7 days. Amplification kinetics were monitored daily by measuring the fluorescence of Thioflavin T (ThT) (Sigma-Aldrich, T3516) using a microplate reader (Thermo Fisher, Varioskan LUX) at excitation/emission wavelengths of 450/485 nm. Following a 7-day incubation, the SAA products were isolated from the reaction mixture. To remove residual Triton X-100, the mixture was added to a 3 kDa MWCO centrifugal filter unit (Millipore, UFC5003) with 15 mL of PBS and centrifuged at 4,000 × g for 30 min at 4°C. The sample was washed eight times using this procedure. To isolate the final product, the purified assemblies were pelleted by centrifugation at 20,000 × g for 30 min. The concentration of remaining monomeric α-synuclein in the supernatant was measured with a BCA assay, and the pelleted PD αSyn preformed fibrils (PD-PFF) were subsequently resuspended in PBS buffer. Prior to use, PD-PFF were sonicated for 30 seconds (0.5-second pulse on/off cycle, 20% amplitude) (53). Aliquots of PD-PFF were stored at – 80°C before use.

### Primary neuronal culture and PD-PFF transduction

Primary cortical neurons were prepared from the cortices of E15.5 humanized *SNCA* mouse embryos. After removal of meninges, cortical tissue was dissociated using 0.25% trypsin-EDTA (Thermo Fisher, 25200056) and triturated into a single-cell suspension, which was then passed through a 40 µm cell strainer (Sigma-Aldrich, CLS431750). Cells were plated onto poly-L-lysine (Sigma-Aldrich, P6407)-coated plates and maintained in neurobasal medium supplemented with B27 (Thermo Fisher, 17504044), 0.5 mM L-glutamine (Thermo Fisher, 25030081), and penicillin/streptomycin (Gibco, 15140163). To inhibit glial proliferation, 5-fluoro-2’-deoxyuridine (1 mM) (Sigma-Aldrich, F0503) was added on day *in vitro* (DIV) 3. On DIV 14, cultures were treated with PD-PFF (54) (10 µg/mL final concentration) or PBS as a vehicle control. Cells were subsequently cultured for an additional 10 days for pathological assays or an additional 14 days for toxicity assays.

### Human iPSC-derived neuron culture and PD-PFF treatment

Cortical glutamatergic neurons were differentiated from the human normal iPSC line iPSC1 as reported (55). iPSCs were stably transfected with a lentiviral vector (56) harboring a doxycycline-inducible Ngn2 transgene, a pro-neural transcription factor widely used for cortical glutaminergic neuron generation from iPSCs (49). Ngn2-driven neuron differentiation was performed following our published protocol (9). To initiate differentiation, the progenitor cells were treated with doxycycline (DOX, Sigma-Aldrich, P6407) for 5 days. Following the induction period, the cells were dissociated and replated onto pre-coated glass coverslips for maturation. The culture medium was then switched to Neurobasal medium (Thermo Fisher, 21103049) supplemented with B27, brain-derived neurotrophic factor (BDNF, PeproTech, 450-02), and glial cell-derived neurotrophic factor (GDNF, PeproTech, 450-10). The neuronal cultures were maintained with a half-medium change performed every three days. The neurons were treated with 10 μg/mL PD-PFF on DIV 14. Samples were subsequently collected at different time points for specific analyses: DIV 16 for the analysis of αSyn mRNA level and total protein level (via immunofluorescence), DIV 28 for the assessment of αSyn-pS129 pathology, and DIV 35 for neurotoxicity assays.

### Immunofluorescence of cultured neurons

Immunofluorescence staining was performed on cultured human cortical neurons and humanized *SNCA* neurons. Briefly, cells were fixed with 4% paraformaldehyde for 15 min, permeabilized, and blocked with 10% goat serum (Thermo Fisher, 10000C) for 1 h at room temperature. Primary antibody incubation was performed overnight at 4 °C using anti-pS129 αSyn (1:1000, Abcam, ab51253), anti-αSyn (1:1000, BD Biosciences, 610787) and anti-NeuN (1:500, Abcam, ab177487). After washing with PBS containing 0.1% Tween 20 (Sigma-Aldrich, P7949), cells were incubated for 1 h at room temperature with appropriate Alexa Fluor-conjugated secondary antibodies (1:1000, Alexa Fluor 488, Thermo Fisher, A10680; Alexa Fluor 568, Thermo Fisher A11004; Alexa Fluor 647, Thermo Fisher, A21244) and Hoechst 33342 (1:5000, Thermo Fisher, 62249) for nuclear counterstaining. Immunofluorescence images were acquired using a Keyence BZ-X710 microscope (Itasca, IL, USA) and a Zeiss LSM880 confocal microscope (Dublin, CA, USA). All images were quantified using the Fiji distribution of ImageJ.

## Acknowledgments

T.M.D. is the Leonard and Madlyn Abramson Professor in Neurodegenerative Diseases. **Funding:** NIH R35GM147498 (JW), Parkinson’s Foundation PF-IMP-1045798 (JW and XM), PF-JFA-1933 (XM), Maryland Stem Cell Research Foundation 2019-MSCRFD-4292 (XM) and 2024-MSCRFD-6394 (XM).

## Author contributions

JW and XM originated the idea of the project. JW and XM designed and led the project and contributed to all aspects of the study; NW, SG and ZT contributed to the experiments, data analysis, and interpretation; NW, SH, ZT, HL, LN, GF and HL contributed to biochemical, cellular, mouse experiments; KG, RK, SZ, AP, TMD, and LSR provided key tools, reagents or critical information or dataset; MY helped designed research and data interpretation; XM and JW wrote the paper. All authors reviewed, edited, and approved the paper.

