## Supplementary Information for "Targeted α-Synuclein mRNA Degradation by PMO-Based RNA-Degrading Chimeras"

**Affiliations:**
[1]Neuroregeneration and Stem Cell Programs, Institute for Cell Engineering, Johns Hopkins University School of Medicine, Baltimore, MD, USA.
[2]Section of Genetic Medicine, Department of Medicine, Biological Sciences Division, University of Chicago, Chicago, IL, USA
[3]Department of Neurology, Johns Hopkins University School of Medicine, Baltimore, MD, USA.
[4]Solomon H. Snyder Department of Neuroscience, Johns Hopkins University School of Medicine; Baltimore, MD, USA.
[5]Department of Physiology, Pharmacology and Therapeutics, Johns Hopkins University School of Medicine; Baltimore, MD, USA.
[6]Institute for NanoBioTechnology, Johns Hopkins University; Baltimore, MD, USA.
[7]Hugo W. Moser Research Institute at Kennedy Krieger; Baltimore, MD, USA.
[8]Department of Materials Science and Engineering, Johns Hopkins University; Baltimore, MD, USA.

†Ning Wang, Shalakha Hegde, Zhichao Tang, Haiqing Liu contributed equally to this work.
φXM and JW contributed equally as senior authors.

**Supplementary Figures**

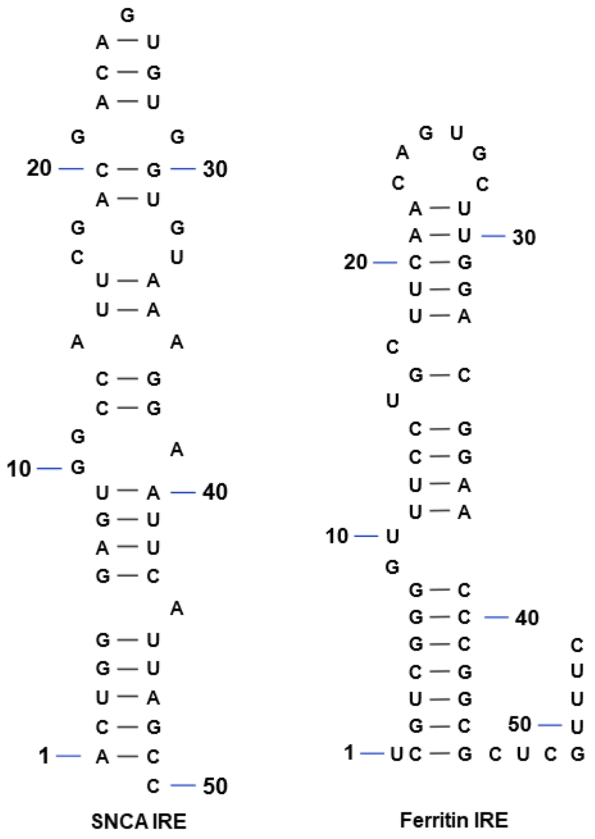


SNCA IRE: 5’ ACUGGGAGUGGCCAUUCGACGACAGUGUGGUGUAAAGGAAUUCAUUAGCC

Ferritin IRE: 5’ UCGUCGGGGUUUCCUGCUUCAACAGUGCUUGGACGGAACCCGGCGCUCGUUUC

**Supplementary Figure 1: Representative secondary structures of IRE.** The stem loop structure differences between the IRE within the 5′ UTR of the *SNCA* transcript (left) and that found in the 5′ UTR of the FTH1 (H-ferritin) transcript (right). The loop regions from each structure are included in the boxes, with a conserved CAGUG motif (highlighted in red), which is a canonical feature of functional IREs and is present in both structures.

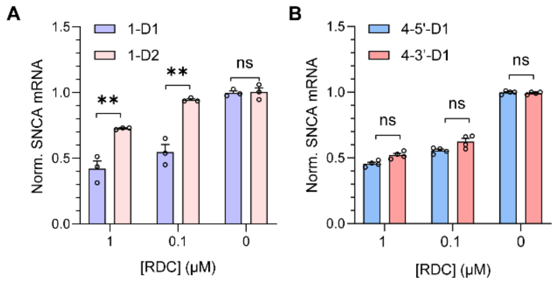


**Supplementary Figure 2: Structural optimization of RDC.** Normalized *SNCA* mRNA expression levels in HEK293T cells measured using RT-qPCR, normalized to the *GAPDH* gene, following 48 h treatment with 1 μM and 0.1 μM of (A) 1-D1 and 1-D2 and (B) 4-5’-D1 and 4-3’-D1. Results are shown as mean ± SEM ($n$ = 4) relative to untreated control. Statistical significance is denoted as follows: * $p < 0.05$, ** $p < 0.01$, ns = not significant ($p > 0.05$), using unpaired two-tailed $t$-tests.

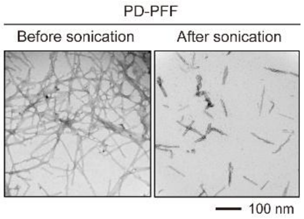


**Supplementary Figure 3: PD-PFF validation.** Typical Transmission electron microscopy images of amplified α-synuclein (αSyn) preformed fibrils of Parkinson disease patient (PD-PFF) before and after sonication. Scale bar: 100 nm.

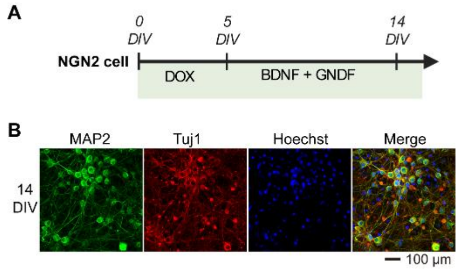


**Supplementary Figure 4: Human cortical neurons validation.** (A) Schematic of the human iPSC culture. (B) Immunofluorescence analysis of MAP2 (green), Tuj1 (red) and Hoechst (blue) in human cortical neurons. Scale bar: 100 µm.

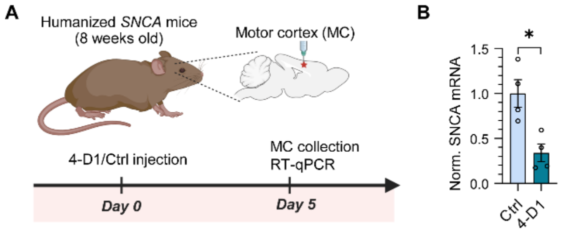


**Supplementary Figure 5: 4-D1 reduced SNCA mRNA level in motor cortex of humanized *SNCA* mice.** (A) Schematic of the experiment design of 4-D1 or Ctrl injection in humanized *SNCA* mice. (B) *SNCA* mRNA level was decreased in 4-D1 injected MC compared to Ctrl group ($n$ = 4). Statistical significance is denoted as follows: * $p < 0.05$, using unpaired two-tailed $t$-tests.

**Supplementary Tables**

Supplementary Table 1: Sequences of PMOs used in the study

| PMO Name | PMO Sequence (5′→3′) | Modification |
|---|---|---|
| *SNCA*_PMO1 | TACACCACACTGTCGTCGAATGGCC | 3′-Alkyne |
| *SNCA*_PMO2 | CGCACCTCACTTCCGCG | 3′-Alkyne |
| *SNCA*_PMO3 | CACACTGTCGTCGAATG | 3′-Alkyne |
| *SNCA*_PMO4 | GAATTCCTTTACACCAC | 3′-Alkyne |
| *SNCA*_PMO5 | GGCCTTTGAAAGTCCTT | 3′-Alkyne |
| *SNCA*_PMO6 | TTGAAGGCAAGGCGTGA | 3′-Alkyne |
| *SNCA*_PMO7 | TGGAAAGGCAGAAGGCTT | 3′-Alkyne |
| *SNCA*_PMO8 | CAGTTCTCCGCTCACGA | 3′-Alkyne |
| *SNCA*_PMO9 | TTCATGAATACATCCAT | 3′-Alkyne |

### Supplementary Methods

#### HEK293T Cell Culture

HEK293T cells (Thermo Fisher, R70007) were cultured in DMEM growth medium (Gibco, 11995040) supplemented with 10% FBS (Cytiva, SH30910.03) and 1% Antibiotic-Antimycotic (Gibco, 15240062) at 37 °C in 5% $CO_2$ atmosphere.

#### Generation of Cas9 expressing cell line

The wild type 293T cells were seeded at 2 × $10^5$ cells/mL in 12-well plates in 1 mL growth medium at 37 °C for 3 h. Cells were transduced with Cas9-expressing lentivirus (Invitrogen, A32064) at a multiplicity of infection (MOI) of 5.0, in the presence of 8 µg/mL polybrene (Sigma-Aldrich, TR-1003-G) to enhance transduction efficiency. At 24 h post-transduction, the culture medium was replaced with fresh growth media. After recovery for 24 h, the transduced cells were then selected in blasticidin (10 µg/mL, Invivogen, ant-bl) for 2 weeks. For stable single clone selection, the cells were diluted in the growth medium containing blasticidin (10 µg/mL) to a final density of 1 cell per 100 µL. The diluted cell suspension was then dispensed to a 96-well plate (100 µL per well). The plate was incubated at 37 °C for 4 weeks. Multiple single-cell colonies were isolated and expanded. To assess Cas9 expression, the expanded cells were lysed in 1× RIPA buffer (Cell Signaling Technology, 9806), and the resulting protein lysates were analyzed by western blotting, as described in the Immunoblotting section of the Methods. Western blot analysis was performed using anti-Cas9 antibody (Cell Signaling Technology, 14697) and γ-tubulin (Sigma-Aldrich, T6557).

#### Generation of RNaseL knockout cell line

Cas9-expressing HEK293T cells were seeded at a density of 3 × $10^5$ cells/ml in 12-well plates with 1 mL of growth medium and incubated at 37 °C for 3 h. Cells were then transduced with a lentiviral LentiArray CRISPR guide RNA (Thermo Fisher, CRISPR ID# 819906; Cat# CRISPR819906_LV), targeting the human RNASEL gene (guide RNA sequence: 5′-UAACGCAGUACAAAUGAGCA). Transductions were performed at a multiplicity of infection (MOI) of 5.0 in the presence of 8 µg/mL polybrene. At 24 h post-transduction, the culture medium was replaced with fresh growth medium. Following an additional 24-h recovery period, transduced cells were subjected to dual antibiotic selection with 2 µg/mL puromycin (InvivoGen, ant-pr-1) and 10 µg/mL blasticidin (InvivoGen, ant-bl) for 2 weeks to enrich for successfully edited cells. For single-cell cloning, the selected population was diluted in selection medium to a final density of 1 cell per 100 µL and seeded into 96-well plates (100 µL per well). Plates were incubated at 37 °C for 4 weeks to allow clonal expansion. Individual colonies were screened for RNase $L^{-/-}$ expression. To assess RNase L expression, the expanded cells were lysed, and the resulting protein lysates in 1× RIPA buffer (Cell Signaling Technology, 9806) were analyzed by western blotting, as described in the Western blotting section of the Methods. Western blot analysis was performed using anti-RNase L antibody (Cell Signaling Technology, 27281) and γ-tubulin.

#### Sanger Sequencing of the RNASEL CRISPR Target Region

Genomic DNA was extracted from CRISPR-edited HEK293T cells and WT 293T cells using the DNeasy Blood & Tissue kit (Qiagen, 69504) according to the manufacturer's instructions. A genomic region flanking the RNase L CRISPR-Cas9 target site was PCR-amplified using primers, RNase L-KO-FW (5′-GCAATTGCTGGAAAGGTGGAG) and RNase L-KO-RV (5′-AAGATTCCTGCTGTCAAGT). PCR amplification was performed using Phusion High-Fidelity DNA Polymerase (Thermo Fisher, F530S) following the manufacturer's protocol. The PCR amplicons were resolved on a 1% agarose gel and the desired DNA bands were excised and purified using Zymoclean Gel DNA Recovery kit (Zymo Research, D4001) according to the manufacturer's instructions. Gel-purified DNA was then submitted for Sanger sequencing using the PCR primers.

Sequencing chromatograms were analyzed using SnapGene Viewer and aligned to the RNase L reference sequence to identify indels.

### Chemistry

Reagents and solvents were purchased from commercial sources (Fisher, Sigma-Aldrich, and Combi-Blocks). The 3'-alkyne-modified PMOs were purchased from Gene Tools (Philomath, OR, USA). D2 was purchased from Wuxi AppTech (Shanghai, China) and used as received. Reactions were tracked by TLC (Silica gel 60 $F_{254}$, Merck) and Agilent 1290 Infinity II HPLC-MS system (Agilent 1290 Infinity II in tandem with LC/MSD IQ). Intermediates and products were purified by a Teledyne ISCO Combi-Flash system using prepacked $SiO_2$ cartridges. NMR spectra were acquired on a Bruker AV400 instrument (400 MHz for $^1$H NMR, 100 MHz for $^{13}$C NMR) or Bruker AV500 instrument (500 MHz for $^1$H NMR, 126 MHz for $^{13}$C NMR). $^{13}$C shifts were obtained with $^1$H decoupling. MestReNova 14.0.1 developed by Mestrelab Research (Santiago de Compostela, Spain) was used for NMR data processing. MS-ESI spectra were recorded on Agilent LC/MSD IQ Mass Detector. HPLC was performed on Agilent 1260 Infinity II PLRP-S C18 (4.6 mm × 150 mm, 5 µm) column and peak detection at 254 nm with UV.

#### Synthesis of D1

#### Ethyl(Z)-5-(4-(2-(2-azidoethoxy)ethoxy)-3-hydroxybenzylidene)-4-oxo-2-(phenylamino)-4,5 dihydrothiophene-3-carboxylate

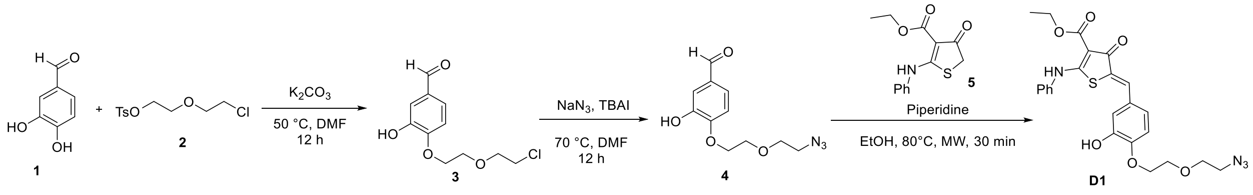


3,4-dihydroxybenzaldehyde 1 (0.1 g, 0.72 mmol) in DMF (2 mL) was added 2-(2-chloroethoxy)ethyl 4-methylbenzenesulfonate 2 (0.19 g, 0.72 mmol) and potassium carbonate (0.1 g, 0.72 mmol). The reaction mixture was stirred at 50 °C for overnight. Added water to the reaction mixture, extracted with diethyl ether. The organic phase was washed with brine solution, dried ($Na_2SO_4$) and concentrated in vacuo to afford compound 3 as a colorless oil (0.15 g, 85%). MS-ESI (*m*/*z*) $[M+1]^+$ 245.00, 247.01.

Compound 3 (0.15 g, 0.56 mmol) in DMF (2 mL) was added sodium azide (0.06 g, 0.83 mmol) and tetrabutyl ammonium iodide (0.23 g, 0.11 mmol). The reaction mixture was stirred at 70 °C for overnight. Added water to the reaction mixture, extracted with diethyl ether. The organic phase was washed with brine solution, dried ($Na_2SO_4$) and concentrated in vacuo. The crude product was purified by silica gel column chromatography (0 – 18% EtOAc in hexanes) to afford compound 4 as a yellow oil (0.08 g, 57%). MS-ESI (*m*/*z*) $[M+1]^+$ 252.08.

$^1$H NMR (400 MHz, Chloroform-*d*) δ 9.88 (s, 1H), 7.48 (d, *J* = 2.0 Hz, 1H), 7.44 (dd, *J* = 8.2, 2.0 Hz, 1H), 7.03 (d, *J* = 8.2 Hz, 1H), 4.37 – 4.29 (m, 2H), 3.98 – 3.88 (m, 2H), 3.83 – 3.75 (m, 2H), 3.50 – 3.42 (m, 2H).

Compound 4 (0.05 g, 0.199 mmol) in ethanol was added ethyl 4-oxo-2-(phenylamino)-4,5-dihydro thiophene-3-carboxylate 5 (0.057 g, 0.218 mmol) and piperidine (3 µL, 0.029 mmol). The reaction mixture was heated under microwave irradiation at 100 °C for 45 min. Evaporated the solvent and added ethanol, heated to 50 °C and filtered and washed with cold ethanol to get D1 as a yellow solid, (0.055 g, 57%). MS-ESI (*m*/*z*) $[M+1]^+$ 497.08.

$^1$H NMR (500 MHz, $CDCl_3$) δ 11.52 (s, 1H), 7.76 (s, 1H), 7.55 – 7.48 (m, 2H), 7.41 (d, *J* = 7.7 Hz, 3H), 7.16 (d, *J* = 2.1 Hz, 1H), 7.06 (dd, *J* = 8.3, 2.2 Hz, 1H), 6.93 (d, *J* = 8.3 Hz, 1H), 4.44 (q, *J* =

7.1 Hz, 2H), 4.25 (dd, *J* = 5.3, 3.7 Hz, 2H), 3.91 – 3.84 (m, 2H), 3.77 (t, *J* = 4.8 Hz, 2H), 3.45 (t, *J* = 4.8 Hz, 2H), 1.47 (t, *J* = 7.0 Hz, 3H).

$^{13}$C NMR (126 MHz, DMSO-$d_6$) δ 177.4, 171.5, 162.3, 142.3, 142.1, 132.4, 126.7, 125.2, 123.8, 123.1, 121.2, 119.4, 114.8, 111.2, 108.9, 65.5, 64.6, 64.3, 55.9, 45.8, 9.7.

**Synthesis of PMO4-D1 RDC**

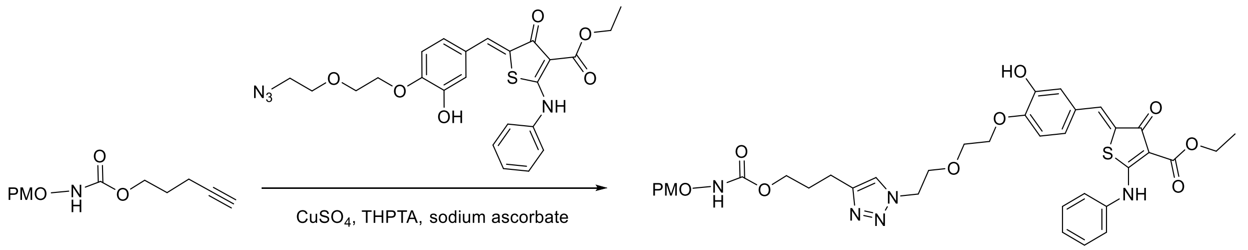


Following the general procedure, using 50 µL *SNCA* PMO4 (1 mM in water), 100 µL *SNCA* PMO4-D1 (0.28 mM in water, measured by Nanodrop) was obtained, with a yield of 56%. MS-ESI(+): 1259.0 [M+5]/5, 1049.1 [M+6] /6, 899.5 [M+7] /7, 787.2 [M+8] /8, 699.9 [M+9]

**Synthesis of PMO4-D2 RDC**

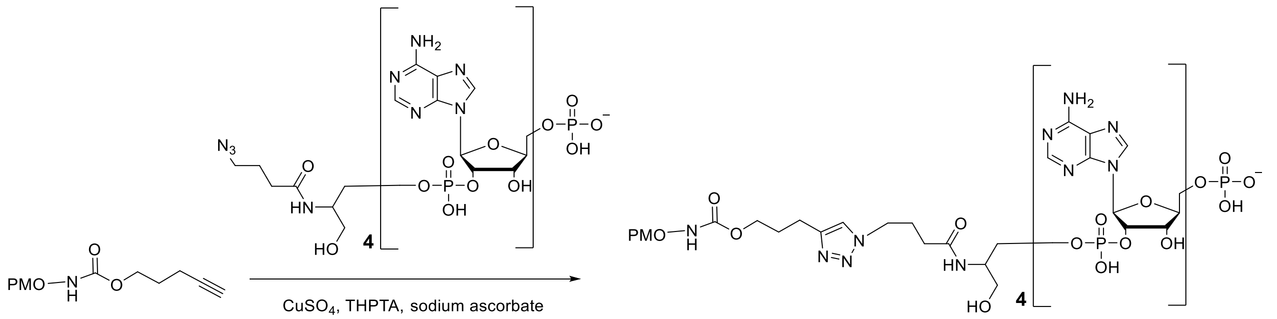


Following the general procedure, using 50 µL *SNCA* PMO4 (1 mM in water), 100 µL *SNCA* PMO4-D2 (0.2 mM in water, measured by Nanodrop) was obtained, yield 40%. MS-ESI(+): 1232.8 [M+6] /6, 1056.7 [M+7] /7, 924.8 [M+8] /8, 822.2 [M+9] /9.

### HPLC analysis of PMO4-D1

**Single Injection Report** — Agilent

| | | | |
|---|---|---|---|
| **Data file:** | D:\CDSProjects\LCMS_Wang\Results\ZT\1_ASO_PMO\Wang_LCMS-2025-08-12 18-35-24-05-00.sirslt\Wang_LCMS-2025-08-12 18-35-24-05-00.dx | | |
| **Sample name:** | ZT-PMO4-RLR_freezedry | | |
| **Description:** | | | |
| **Sample amount:** | 0.000 | **Sample type:** | Sample |
| **Instrument:** | Wang_LCMS | **Location:** | P1-C8 |
| **Injection date:** | 2025-08-12 18:36:28-05:00 | **Injection:** | 1 of 1 |
| **Acq. method:** | 10min_ammonium foramte_column 1_0.5min MS start.amx | **Injection volume:** | 10.000 µL |
| **Analysis method:** | GC_LC Area Percent_DefaultMethod.pmx | **Acq. operator:** | SYSTEM (SYSTEM) |
| **Last changed:** | 2024-11-01 18:29:43-05:00 | | |

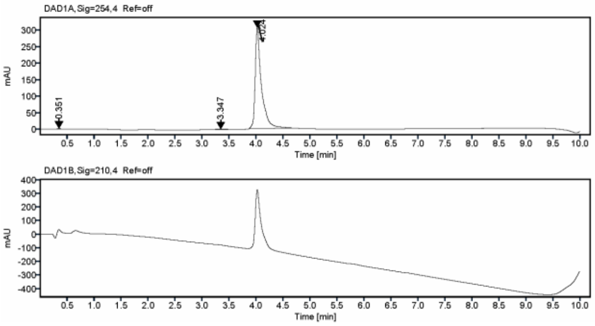


**Signal:** DAD1A,Sig=254,4 Ref=off

| RT [min] | Type | Width [min] | Area | Height | Area% |
|---|---|---|---|---|---|
| 0.351 | MM m | 0.2053 | 12.0093 | 2.8647 | 0.4774 |
| 3.347 | MM m | 0.6182 | 25.4523 | 2.0570 | 1.0117 |
| 4.024 | MM m | 0.8439 | 2478.2128 | 306.3064 | 98.5109 |
| | | **Sum** | **2515.6745** | | |

D:\CDSProjects\LCMS_Wang\Report Templates\ZT_purity.rdl [Rev. 14.0]  Printed: 2025-09-04 18:35:37-05:00  Page 1 of 1

### Stereotaxic injection of 4-D1 or unconjugated PMO4 (Ctrl)

For intracortical injections, 8-week-old humanized *SNCA* mice were deeply anesthetized with a ketamine mixture (100 mg/kg, i.p.) and secured in a stereotaxic apparatus. After exposing the skull, a small burr hole was drilled over the target location to allow for needle access. Using a 10 µL Hamilton syringe, 2 µL of 5 µM 4-D1 or an equivalent volume of Ctrl was unilaterally injected into the motor cortex at the following coordinates relative to bregma: AP: +0.5 mm, ML: +1.0 mm, DV: -0.5 mm. The infusion was performed at a controlled rate of 0.2 µL/min to minimize tissue disruption. To ensure complete diffusion and prevent backflow, the needle remained in place for 10 minutes post-injection before being withdrawn slowly. The incision was then sutured, and the mice received postoperative care, including analgesics for pain relief and monitoring during recovery. 5 days after injection, the mice were humanely euthanized for brain tissue collection and subsequent mRNA analysis.

**αSyn purification**

Recombinant human α-synuclein (αSyn) monomer was purified as previously described. Briefly, the pRK172-αSyn plasmid was transformed into BL21 (DE3) E. coli for protein expression. Following large-scale culture, the expression of αSyn was induced with the addition of isopropyl β-D-1-thiogalactopyranoside (IPTG). The bacterial pellets were then collected by centrifugation and lysed via osmotic shock to release the intracellular proteins. The resulting supernatant, containing the soluble αSyn fraction, was clarified and dialyzed overnight against the appropriate buffer. Monomeric αSyn was then purified from the dialysate by Fast Protein Liquid Chromatography (FPLC) using an anion-exchange column, which separates proteins based on their net charge. The purity of the collected fractions was confirmed by SDS-PAGE, and protein concentration was measured. As a final critical step, bacterial endotoxins were removed using a ToxinEraser Endotoxin Removal Kit (GenScript, L00338) to ensure the protein preparation would not elicit an inflammatory response in subsequent cellular or animal experiments.

**Transmission electron microscopy (TEM) measurements**

For morphological analysis, we used transmission electron microscopy (TEM) with a negative staining protocol to visualize the fibrillar structures. First, a small droplet of the amplified α-synuclein pre-formed fibrils from a Parkinson's disease patient (PD-PFF) was carefully applied to the surface of a 400-mesh carbon-coated copper grid (Electron Microscopy Sciences, CF400-Cu-50), which provides an ideal electron-transparent support film. After a 2-minute adsorption period, the grids were washed with double-distilled water to remove unbound fibrils and residual buffer salts that could otherwise crystallize and obscure the image. To enhance contrast, the grids were then negatively stained with 2% uranyl acetate (Electron Microscopy Sciences, 22400) for 1 minute. This heavy metal salt solution pools around the lighter protein fibrils, embedding them in an electron-dense background that effectively highlights their shape and structure. After staining, the excess liquid was carefully wicked away using filter paper, and the grids were allowed to air-dry completely overnight. Finally, grids were imaged using a Hitachi H7600 TEM at an acceleration voltage of 80 kV to confirm the presence of the characteristic filamentous morphology. Sample preparation and concentration were kept consistent across all experiments to ensure reproducibility.

**Quantification and Statistical Analysis**

All statistical analyses were conducted using GraphPad Prism software (Version 8), a standard tool for biological data analysis. The specific statistical test was chosen based on the experimental design; a two-tailed unpaired t-test was used for direct comparisons between two groups, whereas a one-way analysis of variance (ANOVA) was employed for comparisons across three or more groups. When a significant difference was detected by ANOVA, the indicated post-hoc test (e.g., Tukey's or Dunnett's multiple comparisons test) was performed to identify which specific group means were different. Quantitative data are expressed as the mean ± standard error of the mean (SEM) or, in some cases, shown as violin plots to better visualize the distribution and include all individual data points. To ensure the robustness and reproducibility of our findings, all experiments were performed with at least three independent biological replicates. Similarly, any morphological images shown are representative of at least three independent experiments that yielded similar outcomes. For all statistical tests, a p-value of less than 0.05 was considered the threshold for statistical significance.

**Abbreviations**

AAV: adeno-associated virus

αSyn: alpha synuclein

ASO: antisense oligonucleotide

BDNF: brain-derived neurotrophic factor

BSA: bovine serum albumin

DOX: doxycycline

GDNF: glial cell-derived neurotrophic factor

iPSCs: Atoh1-transduced induced pluripotent stem cells

MSA: Multiple System Atrophy

PD: Parkinson’s disease

PFF: preformed fibrils

PMO: phosphorodiamidate morpholino oligonucleotide

pS129: phosphorylated at serine 129

RDC: RNA-degrading chimeras

RISC: RNA-induced silencing complex

RLR: RNase L recruiter

RNAi:  RNA interference

SAA: seeding amplification assay

ShRNA: short hairpin RNAs

SiRNA: small interfering RNAs

TEM: transmission electron microscopy

ThT: Thioflavin T

UTR: untranslated region
